# Cell survival enabled by leakage of a labile metabolic intermediate

**DOI:** 10.1101/2022.05.20.492833

**Authors:** Encarnación Medina-Carmona, Luis I. Gutierrez-Rus, Fadia Manssour-Triedo, Matilda S. Newton, Gloria Gamiz-Arco, Antonio J. Mota, Pablo Reiné, Juan Manuel Cuerva, Mariano Ortega-Muñoz, Eduardo Andrés-León, Jose Luis Ortega-Roldan, Burckhard Seelig, Beatriz Ibarra-Molero, Jose M. Sanchez-Ruiz

## Abstract

Many metabolic pathways are of ancient origin and have evolved over long periods of time (Noda-Garcia et al., 2018). Yet, new pathways can also emerge in short time scales in response, for instance, to the presence of anthropogenic chemicals in the environment (Copley, 2009). Models of metabolic pathway emergence and evolution often emphasize the acquisition of new reactions through horizontal gene transfer and promiscuous enzyme functionalities (Pál et al., 2005; Schulenburg & Miller, 2014; Copley, 2015; Noda-Garcia et al., 2018; Peracchi, 2018). A fundamentally different mechanism of metabolic innovation is revealed by the evolutionary repair experiments reported here. A block in the proline biosynthetic pathway that compromises cell survival is efficiently rescued by many single mutations (12 at least) in the gene of glutamine synthetase. The mutations cause the leakage to the intracellular milieu of a sequestered phosphorylated intermediate common to the biosynthetic pathways of proline and glutamine, thus generating a new route to proline. Metabolic intermediates may undergo a variety of chemical and enzymatic transformations, but are typically protected as shielded reaction intermediates or through channeling in multi-enzyme complexes and metabolons (Srere, 1987; Huang et al., 2001; Grunwald, 2018; Pareek et al., 2021). Our results show that intermediate leakage can readily occur and contribute to organismal adaptation. Enhanced availability of reactive molecules may enable the generation of new biochemical pathways. We therefore anticipate applications of mutation-induced leakage in metabolic engineering.

---

Determining the molecular mechanisms responsible for the emergence and evolution of metabolic pathways is important for our understanding of how life has evolved. Furthermore, detailed information about these mechanisms may suggest new tools for synthetic biology and metabolic engineering. Re-enacting the natural evolution of metabolic pathways in the lab is certainly challenging. However, fundamental mechanisms of metabolic innovation can be revealed through experiments with cells in which a crucial enzyme has been deleted. The deletion blocks a metabolic pathway, makes the cells auxotrophic for its final product and allows evolutionary experiments on the restauration of prototrophy to be performed in the lab. A number of such evolutionary repair experiments have provided support for the proposed role of enzyme promiscuity in the emergence of new biochemical pathways (Patrick et al., 2007; McLoughlin & Copley, 2008; Kim et al., 2010; Digianantonio & Hetch, 2016; Digianantonio et al., 2017; Kim et al., 2019). Note that we are using the term “enzyme promiscuity” in a general sense, *i*.*e*., including the enzyme capability to catalyze a given chemical reaction with different substrates (substrate scope), as well as the capability to catalyze different reactions (catalytic promiscuity). Certainly, the low-level promiscuous activities displayed by many enzymes often have no physiological role (Khersonsky & Tawfik, 2010; Copley, 2015). Yet, their enhancement upon mutation may allow the organism to meet new challenges by patching novel pathways (Kim et al., 2019).

Here, we report evolutionary repair experiments on *E. coli* cells in which an enzyme crucial for the biosynthesis of proline has been deleted. Remarkably, we find reversion to proline prototrophy through a mechanism that does not involve the recruitment of a low-level promiscuous activity, but rather the leakage to the intracellular milieu of a shielded metabolic intermediate.

In most organisms, proline is synthesized from glutamate through a pathway that involves four steps (Csonka & Leisinger, 2013; Fichman et al., 2015) (Fig. 1). The first step is the phosphorylation of glutamate to γ-glutamyl phosphate, which is followed by reduction to γ-glutamyl semialdehyde, cyclization to Δ^1^-pyrroline-5-carboxylate and further reduction to proline. While the cyclization step is spontaneous, the first, second and fourth steps are catalyzed by the enzymes γ-glutamyl kinase, γ-glutamyl phosphate reductase and Δ^1^-pyrroline-5-carboxylate reductase, respectively. In *E. coli*, the genes encoding these enzymes are known as *proB, proA* and *proC*, respectively (see Fig. 1).

**Fig. 1.**
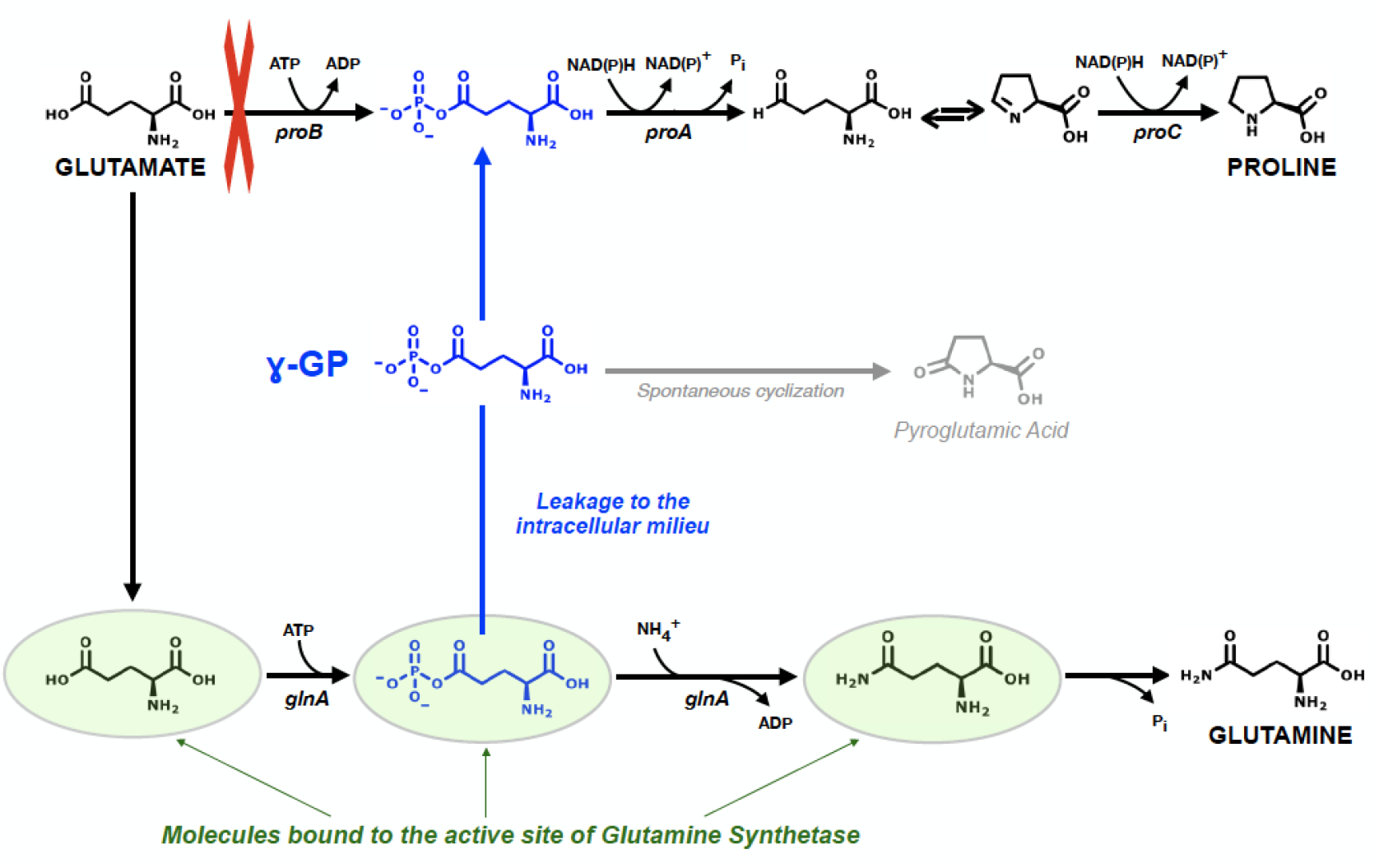
Biosynthetic pathways of proline and glutamine. A block in the proline biosynthetic pathway (red cross: deletion of the enzyme encoded by the *proB* gene) is rescued by mutations in the gene encoding glutamine synthetase that cause the leakage of γ-glutamyl phosphate (γ-GP), a high-energy phosphorylated compound. The leaked γ-GP can either undergo spontaneous cyclization to yield 5-oxopyrrolidine-2-carboxylic acid (pyroglutamic acid) or diffuse to become substrate of γ-glutamyl phosphate reductase, the enzyme encoded by the *proA* gene. See text for details. Note that glutamine biosynthesis from glutamate occurs at the active site of glutamine synthetase. To show this fact, glutamine synthetase is represented by a green oval enclosing the molecules bound to its active site. Leakage of γ-glutamyl phosphate from the active site of the synthetase and its transfer to the active site of γ-glutamyl phosphate reductase (the product of the *proA* gene) is highlighted in blue.

Here, we have studied a *ΔproB* strain from the Keio collection (Baba et al., 2006). This *E. coli* Keio knockout cannot produce γ-glutamyl kinase, the first enzyme of the proline biosynthetic pathway, and, as a result, cannot grow in the absence of proline in the culture medium. However, upon plating of the knockout strain onto minimal medium, we found that a small number of colonies were visually apparent after only a few days. Specifically, we prepared several hundred plates, which we examined for four days. Colonies appeared during days 2, 3 and 4 (Extended Data Fig. 1) and most of the colonies, up to a total of 30, showed robust growth and gave rise to many new colonies upon re-plating onto minimal medium (Extended Data Fig. 2).

Early studies (Berg & Rossi, 1974) reported that some proline auxotrophs blocked at the first or second steps in the proline biosynthetic pathway could revert to prototrophy as a result of mutations in the arginine biosynthetic pathway. Briefly, impaired activity of the enzyme encoded by the *argD* gene leads to accumulation of the substrate of this enzyme, N-acetylglutamyl semialdehyde, which, with the assistance of a promiscuous activity of the enzyme encoded by the *argE* gene, is transformed into γ-glutamyl semialdehyde, thus bypassing the block in proline biosynthesis.

Surprisingly, sequencing of our rescued *E. coli* cells did not detect any mutations in the enzymes involved in arginine biosynthesis, but showed that most of our rescues had single mutations in the gene for glutamine synthetase, *glnA*. We performed Illumina whole genome sequencing on the seven rescues that appeared first (*i*.*e*., on the second day after plating). We found no mutations in 2 of them, suggesting that, in these cases, rescue is linked to alterations that cannot be easily detected by random fragment sequencing (gene duplications or transposon insertions, for instance). On the other hand, we found single mutations in the Illumina whole-genome sequencing of five of the rescues, but the mutations appeared solely in the gene of glutamine synthetase. Furthermore, 19 (out of 21) of the rescues that appeared on the third and fourth days after plating had single mutations in the gene of glutamine synthetase, as shown by Sanger sequencing. In total, we identified 15 mutations in the glutamine synthetase gene of the *E. coli* cells that reversed to proline prototrophy. Some of those mutations appeared in up to three of the independently generated rescues. In order to test whether the mutations are sufficient for rescue, we complemented the *ΔproB* knockout cells with a plasmid expressing the mutated variants of glutamine synthetase, as well as the wild type enzyme as control. We observed efficient growth in the absence of proline for 12 of the identified mutations, while no growth was observed upon complementation with the wild type enzyme (Extended Data Fig. 3).

Glutamine synthetase, a central enzyme in nitrogen metabolism, catalyzes the ATP-dependent synthesis of glutamine from glutamate and ammonia (Eisenberg et al., 2000; Stadtman, 2013). The enzyme is a dodecamer of identical subunits (Fig. 2a) with active sites formed by residues from two monomers (Almassy et al., 1986). The reaction occurs via two steps (see Fig. 1): the first step is the formation of the intermediate γ-glutamyl phosphate, which subsequently reacts with ammonia in a second step to yield glutamine with the concomitant release of phosphate (Eisenberg et al., 2000). Note that the γ-glutamyl phosphate intermediate in the catalytic cycle of glutamine synthetase is the same as the product of the reaction catalyzed by γ-glutamyl kinase, the first enzyme in the proline biosynthetic pathway which is deleted in the *ΔproB* knockout strain. Therefore, it seems reasonable to hypothesize that the mutations responsible for the restoration of proline prototrophy in the *ΔproB* knockouts cause the leakage of the intermediate from the active site of the synthetase to the intracellular milieu and that diffusion then allows the intermediate to become the substrate of γ-glutamyl phosphate reductase (Fig. 1). Indeed, the rescuing mutations define a clear structural pattern, being in all cases at the interaction surface between two glutamine synthetase monomers and close to the active site (Fig. 2b).

**Fig. 2.**
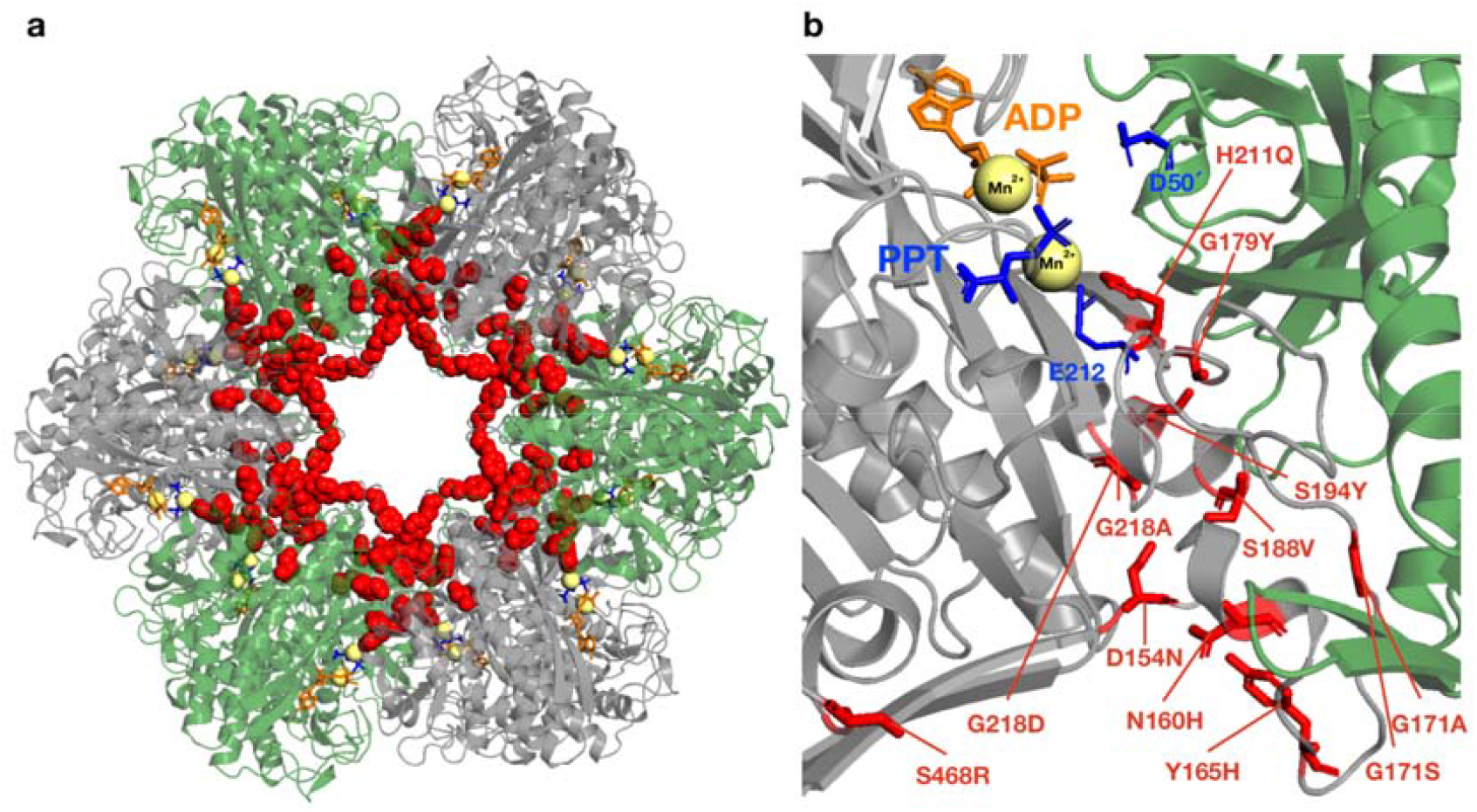
Location of the mutations that rescue proline biosynthesis in the structure of glutamine synthetase. **(a)** Bacterial glutamine synthetases form dodecamers of identical subunits with two face-to-face rings (only one ring is shown), with the active sites located at the interfaces between the subunits at each hexameric ring (Eisenberg, 2000). The mutations that rescue *ΔproB* (original residues shown with red spheres) accumulate at the interfaces between subunits. For clarity, neighboring subunits are shown in different color (green and grey) **(b)** The interface between two subunits showing the active site region. Bound manganese cations, ADP and the inhibitor phosphinothricin (PPT), a substrate analog, are shown. The carboxylic acid residues D50’ and E212 are highlighted to indicate the approximate location of the ammonia binding site. Red labels with original residues also in red identify the 12 positions for which rescuing mutations were found. The structure shown in (a) and (b) a corresponds to the glutamine synthetase from *Salmonella typhimurium* (PDB ID: 1FPY), which has 97.9% sequence identity with the *E. coli* enzyme. The residues at the positions labeled in red are identical in both enzymes.

The leakage hypothesis expounded in the preceding paragraph would seemingly face, however, a serious problem: γ-glutamyl phosphate is a high-energy phosphorylated compound that is extremely unstable and readily cyclizes when exposed to water yielding 5-oxopyrrolidine-2-carboxylate (Csonka & Leisinger, 2013; Fichman et al., 2015). Hence, it is generally assumed that γ-glutamyl phosphate can only exist as an enzyme-bound intermediate, as, for instance, in the catalytic cycle of glutamine synthetase. In the case of the proline biosynthetic pathway in *E. coli*, the enzymes responsible for the first and second steps, γ-glutamyl kinase and γ-glutamyl phosphate reductase, are believed to form a complex *in vivo*, thus allowing the channeling of the labile γ-glutamyl phosphate directly from the kinase active site to the reductase active site and preventing undesirable side reactions (Csonka & Leisinger, 2013; Fichman et al., 2015). This notion is supported by the fact that the orthologs of these enzymes in animals and vascular plants are fused to form a single bifunctional enzyme (Csonka & Leisinger, 2013; Fichman et al., 2015).

However, the experiments and calculations we describe below support that, despite its extreme lability, γ-glutamyl phosphate does indeed leak from the active site of glutamine synthetase variants to the intracellular milieu and diffuse to reach γ-glutamyl phosphate reductase, thus bypassing the block in protein biosynthesis generated by the deletion of γ-glutamyl kinase.

Catalytic features of glutamine synthetase are subject to complex modulation related, among other factors, to adenylation-deadenylation and the existence of taut and relaxed conformations (Stadman, 2013). Here, however, we are only concerned with the possibility of leakage of γ-glutamyl phosphate from the catalytic cycle of the enzyme and this can be easily demonstrated through the detection of 5-oxopyrrolidine-2-carboxylate (pyroglutamic acid), the product resulting from γ-glutamyl phosphate cyclization when released from the synthetase active site to water (Fig. 3a). We have thus prepared the wild-type enzyme and its single-mutant variants following the same procedure and we have studied *in vitro* the catalytic reaction under the same solvent conditions. Both NMR and mass spectrometry detected that all of the mutations responsible for the restoration of proline prototrophy cause an increase in pyroglutamic acid consistent with leakage of the γ-glutamyl phosphate intermediate (Figs. 3b and 3c). As was to be expected, there is a substantial trade-off between the leakage of the intermediate and the normal activity of glutamine synthetase (compare the two plots in Fig. 3b). That is, as the leakage of the intermediate increases, the amount of glutamine produced by the enzyme decreases. Still, the production of glutamine is not fully impaired and the mutations that rescue proline biosynthesis should not completely block glutamine biosynthesis.

**Fig. 3.**
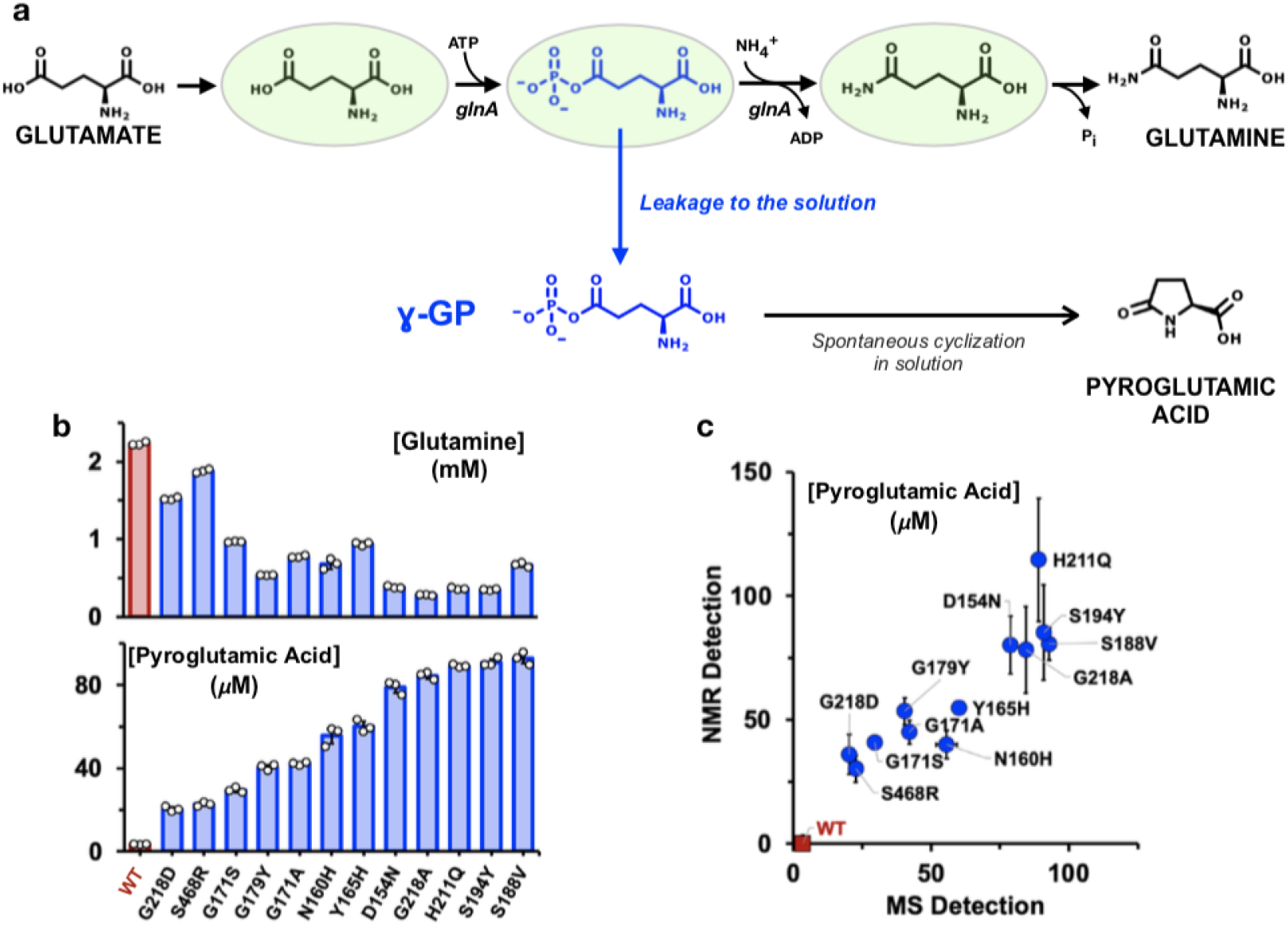
Mutation-induced leakage of γ-glutamyl phosphate (γ-GP) from glutamine synthetase. **(a)** Leakage of γ-GP from the enzyme to the bulk solution can be demonstrated through the detection of the product of its spontaneous cyclization, pyroglutamic acid. Note that glutamine biosynthesis from glutamate occurs at the active site of glutamine synthetase. To show this fact, glutamine synthetase is represented by a green oval enclosing the molecules bound to its active site (see Fig. 1). **(b)** To solutions of 4 mM glutamate and 25 mM in NH_4_Cl at pH 7, variants of glutamine synthetase were added to a final concentration of 0.5 μM. The solutions were analyzed by mass spectroscopy to detect pyroglutamic acid and the glutamine after 12 hours of reaction at 25 °C (see Methods for details). The experiment was performed with wild type glutamine synthetase and 12 single-mutant variants responsible for the rescue of proline biosynthesis (Fig. 2). Values for three biological replicates are shown with open circles. The plots shown reveal the mutation-induced leakage and the trade-off with the normal activity of glutamine synthetase, i.e. efficient leakage implies impaired conversion of glutamate to glutamine. (c) The experiment in (b) was repeated with 3 mM glutamate and detection of pyroglutamic acid by NMR. The plot shows an excellent congruence of the data obtained from NMR with the data obtained from mass spectrometry. The data shown in (b) and (c) are the average of three independent measurements in each case. Therefore, for each variant, the mutation-induced leakage is supported by 6 independent measurements using two different experimental methodologies and at least two different protein batches. Average values and standard deviations are shown.

Mass spectrometry detection of pyroglutamic acid can also be used to explore the leakage of γ-glutamyl phosphate *in vivo*. Several processes can generate pyroglutamic acid in living cells (Kumar & Bachhawat, 2012). In order to probe specifically its formation from glutamate activation, we let several of the original rescues grow in minimal medium to an optical density of 0.7, added isotopically labeled glutamate to a 1 mM concentration and, after ten minutes, used mass spectrometry to detect the corresponding isotopically-labeled pyroglutamic acid. As a control experiment, we used the same protocol with *ΔproB* knockout cells, but supplementing the culture medium with 50 μM proline to enable growth. We found significantly higher levels of pyroglutamic acid in the rescued cells, as compared with the control (Extended Data Fig. 4). Glutamate can yield pyroglutamic acid through a non-enzymatic process (Kumar & Bachhawat, 2012), but this is expected to be slow and occur with the same rate in the rescues and the control. Formation of pyroglutamic acid from the γ-glutamyl phosphate produced by γ-glutamyl kinase is not possible in these cells, since the encoding gene, *proB*, has been deleted. We conclude, therefore, that the enhanced levels of pyroglutamic acid in the cells that rescue proline auxotrophy (Extended Data Fig. 4) reflect the leakage of γ-glutamyl phosphate from the glutamine synthetase variants.

Of course, for leakage to be effective in restoring proline prototrophy, a significant fraction of the γ-glutamyl phosphate leaked from the glutamine synthetase must be able to reach the γ-glutamyl phosphate reductase before undergoing cyclization. Transfer of γ-glutamyl phosphate between the two enzymes can be explored *in vitro* through an obvious modification of the activity assay for γ-glutamyl phosphate reductase (Smith et al., 1984). Since, γ-glutamyl phosphate is a highly unstable molecule, it is supplied in this assay by the previous enzyme of the proline biosynthetic pathway, γ-glutamyl kinase (the product of the *proB* gene: see Fig.1). We have modified this coupled assay by replacing the kinase with one of our glutamine synthetase variants (Fig. 4a). In this way, activity is only observed if the synthetase releases a sufficient amount of γ-glutamyl phosphate that this can reach the reductase before undergoing cyclization. Indeed, reductase activity is observed in these coupled assays (Fig. 4b and Extended Data Fig. 5) for all of the 12 variants of glutamine synthetase that we found to efficiently restore proline prototrophy. In contrast, no reductase activity is detected when using wild type glutamine synthetase. These *in vitro* coupled assays demonstrated that γ-glutamyl phosphate can diffuse from the synthetase variants to the reductase. We propose that the same scenario applies *in vivo*. Supporting this idea, theoretical calculations of the energy barrier for the cyclization of the phosphorylated intermediate suggest a time scale for γ-glutamyl phosphate cyclization in solution within a range of tens of milliseconds to seconds (Extended Data Figs. 6 and 7), while the the diffusion of molecules across an *E. coli* cell is expected is expected to occur in a shorter time scale (Philips et al., 2009) (see legends to Extended Data Figs. 6 and 7 for details). Therefore, within an *E. coli* cell, a substantial fraction of the molecules leaked by the glutamine synthetase variants should be able to reach the γ-glutamyl phosphate reductase before being lost to cyclization. Interestingly, the reductase activity values determined in the coupled assay with the glutamine synthetase variants match qualitatively the general trend observed for the efficiency of the different mutations to restore proline prototrophy (Extended Data Fig. 8).

**Fig. 4.**
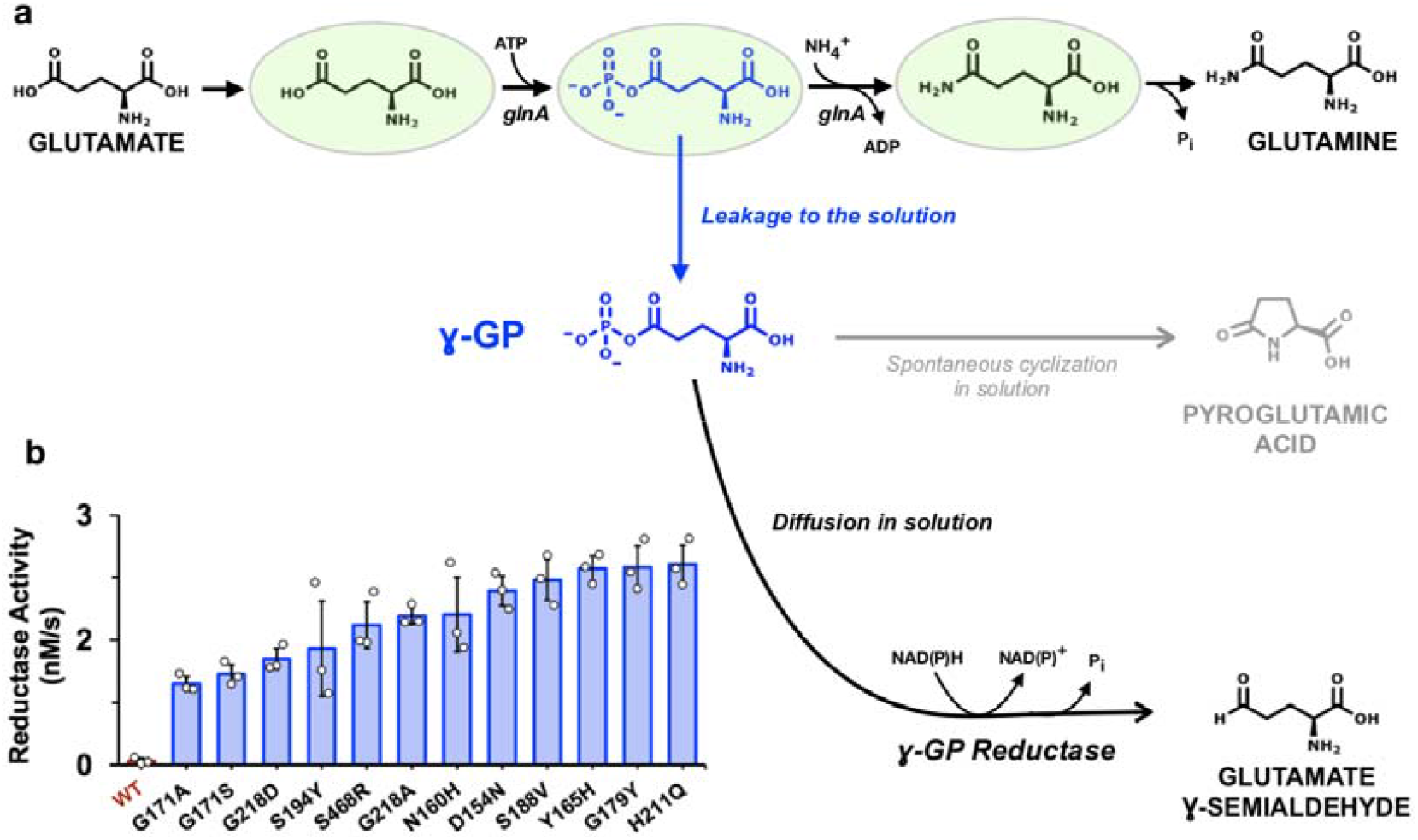
Diffusion of γ-glutamyl phosphate (γ-GP) from glutamine synthetase to γ-GP reductase. γ-GP leaked from glutamine synthetase was detected via a coupled assay with γ-GP reductase. **(a)** The γ-GP leaked from the glutamine synthase can diffuse and be processed by the γ-GP reductase, despite diffusion competing with the spontaneous cyclization of γ-GP. This coupled *in vitro* assay simulates in solution the molecular process responsible for the rescue of a block in proline biosynthesis by γ-GP leakage from glutamine biosynthesis (Fig. 1). Note that glutamine biosynthesis from glutamate occurs at the active site of glutamine synthetase. To show this fact, glutamine synthetase is represented by a green oval enclosing the molecules bound to its active site (see Fig. 1). **(b)** The assay of γ-GP reductase coupled with glutamine synthetase was performed as described in Methods using wild-type synthetase and the single-mutant variants responsible for the rescue of proline biosynthesis (Fig. 2). All variants of glutamine synthetase lead to activity values much higher than the wild-type level, which was undetectable. The values shown are the average of at least three biological replicates (Extended Data Fig. 4) and the original replicate values are also shown with open circles. Error bars are standard deviations.

Possibly the most remarkable aspect of the experimental results reported here is the high proficiency of intermediate leakage at rescuing a block in an essential biochemical pathway. Not only reversion to proline prototrophy is realized by a large number (12 at least) of different single mutations, but also the rescued cells in a culture medium lacking proline show robust growth immediately (Extended Data Figs. 2 and 3). Yet, this surprising proficiency can be convincingly rationalized. At a fundamental level, leakage does not involve the generation of a new molecular functionality, but the impairment of a previous evolutionary adaptation, the sequestering of an intermediate, and damaging a highly-optimized feature can obviously be achieved in many different ways. In general, for instance, mutations that weaken the interactions of the bound intermediate with the protein moiety or mutations that decrease the rate of the enzymatic transformation of the intermediate can enhance the likelihood that the intermediate is released to the intracellular milieu. Certainly, mutations that bring about leakage will also decrease the normal activity of glutamine synthetase thus leading to a trade-off. However, this would be immediately corrected to some substantial extent by the regulation mechanisms of the glutamine biosynthesis pathway. Furthermore, since some of these regulation mechanisms work at the level of transcription (Stadman, 2013), they may increase the expression levels of glutamine synthetase and consequently the flow through the new pathway generated by leakage. In general, it appears that leakage of one intermediate may occur without fatal consequences for the cell.

Sequestration of intermediate chemical species in metabolism is not restricted, of course, to the specific case of the catalytic cycle of glutamine synthetase but occurs extensively and in a variety of molecular scenarios (Srere, 1987; Huang et al., 2001; Grunwald, 2018; Pareek et al., 2021). Reaction intermediates in the cycles of enzymes catalyzing multi-step reactions remain bound in the active site and are shielded from interactions with other molecules present in the intracellular milieu. In some poly-functional enzymes, intermediates that are produced in one active site are transferred to a second active site through an intermolecular tunnel, thus preventing their release to the intracellular milieu. Enzymes that catalyze successive steps in a metabolic pathway often form a multi-enzyme complex in such a way that the intermediate (product of the first step and substrate of the second) is directly channeled between the two enzymes. Cluster channeling of intermediates occurs in metabolons that are temporary associations of sequential metabolic enzymes in which multiple copies of each enzyme are present.

Extensive sequestration is obviously required for metabolic efficiency and widespread leakage of metabolic intermediates to the intracellular milieu will likely result in cell death. On the other hand, our results support that leakage of one given intermediate may readily occur without fatal consequences for the organism and that it may even contribute to organismal adaptation in some scenarios. About 20% of metabolic intermediates have several uses in the cell (Srere, 1987) and rescue of a block in an essential route by leakage of a sequestered intermediate from a different route, as demonstrated here with the proline and glutamine pathways, could plausibly occur in many other instances. Furthermore, metabolic intermediates are often intrinsically reactive species that not only may become alternative substrates for promiscuous enzymes (Noda-Garcia et al., 2018), but that may also undergo various non-enzymatic chemical transformations (Lerma-Ortiz et al., 2016). As a result, leakage of one intermediate upon mutation could facilitate its transformation, enzymatic or non-enzymatic, into different chemical species. Perhaps, some of the new molecules thus generated will be toxic and will need to be neutralized by the metabolite damage-control system of the cell (Crécy-Lagard et al., 2018). Yet, some others could trigger, perhaps with the assistance of promiscuous enzyme functionalities, the generation of new biochemical pathways that allow the organism to meet specific challenges. Additional work will be required to assess the extent to which this leakage mechanism has contributed to the diversity of the products of secondary metabolism or, more generally, to metabolic pathway evolution by generating new metabolites. On the other hand, it is clear that the leakage mechanism can be immediately used as a tool in synthetic biology for metabolic engineering. The design of pathways for the production of valuable products (Lee et al., 2018; Cravens et al., 2019) may obviously benefit from the use of highly reactive molecules as precursors. Certainly, highly reactive molecules are most likely to be sequestered *in vivo*, but this works demonstrated that their availability can be engineered, as leakage-inducing mutations can easily be found by *in vivo* or *in vitro* screening for variants of the crucial proteins involved in sequestration along the general lines of the experiments described here.

## Methods

### Selection and identification of spontaneous proline auxotrophy rescues

*E. coli* BW25113 ΔproB760::kan (JW0232-1) was obtained from the Keio collection (Baba et al., 2006). Aliquots of frozen stocks of the ΔproB knockout strain were streaked onto 240 plates containing M9–glucose medium [agar (2% w/v), M9 salts (1x), glucose (0.4%, w/v), MgSO4 (2 mM), CaCl2 (100 μM)] supplemented with Kanamycin (Kan; 30μg·Ml^-1^). Plates were incubated at 37 °C for up to 4 days, and colony formation was monitored. The rescuing activity of each individual clone was verified by restreaking onto new M9-glucose-Kan plates up to 3 times. Genomic DNA was extracted from isolated colonies for each of the rescue clones with the bacterial genomic DNA isolation kit from Norgen (CAT#17900) and quantified using a Biodrop μLite+.

### Genomic DNA sequencing

Of the total number of clones, DNA from 9 plate samples were sequenced: two control samples (*E. coli* BW25113 ΔproB760:kan (JW0232-1)) and seven samples from rescue clones obtained as explained in the previous section.

These were sequenced on a Novaseq 6000, using an S2 cartridge generating on average 8,392,966 × 150 nt paired reads.

For the remaining gDNA samples from rescue clones, the glnA gene was amplified by PCR using the PfuUltra high-fidelity DNA polymerase AD from Agilent (CAT#600385) and the following specific primers: glnA.for (5’-AGA CGC TGT AGT ACA GCT CAA ACT C) and glnA.rev (5’-ATG TCC GCT GAA CAC GTA CTG ACG ATG). Amplified products were sequenced by Sanger method using specific primers to glnA gene in order to identify the rescue mutations.

### Bioinformatics analyses

Paired files obtained after sequencing were evaluated using FASTQC software to identify possible errors, such as low-quality sequences, accumulation of adapter sequences or unidentified nucleotides (Andrews, 2010).

After this quality control filtering, we observed that 99.1% of the reads had an average quality value higher than Q30, and there was no accumulation of adapters, neither repeated sequences nor a high percentage of Ns. After selecting, by cutadapt (Martin, 2011), the sequences with an average quality value of at least Q30, Snippy was used to detect variants in the sequenced DNA (Seemann, 2015). Briefly, Burrow Wheeler Alignment (Li and Durbin, 2009) was used to align the reads against the reference genome (GCA_000750555 from GeneBank). Samtools (Li et al., 2009) was used to sort, mark and remove duplicate sequences. Once aligned and sorted, depth was calculated to estimate the number of reads that covers each nucleotide, on average we obtained 412X. Subsequently, variant calling was performed using the default parameters by Freebayes (Garrison and Marth, 2012). Of note is that in all samples in which the whole genome was sequenced, including the two controls, a mutation in the cpxA gene (342C>A nucleotide change coding for the Asn114Lys aminoacid change) appeared when compared to the published reference sequence. Since, however, this mutation also appeared in the controls (the non-rescued *ΔproB* knockouts), it is clear that it is not related to reversion to prototrophy.

### Protein mutagenesis and expression

The coding sequence for *E*.*coli* glutamine synthetase (GS) was inserted into the Kpn1/Xho1-digested pET-45b(+) expression vector (GenScript Biotech, Netherlands), containing a hexahistidine tag at its N-terminus and ampicillin resistance gene on the plasmid. Single-mutant variants of glutamine synthetase were prepared by site-directed mutagenesis using the QuikChange Lightning kit (Agilent technologies, USA). The gene coding for γ-glutamyl phosphate reductase was ligated into the pHTP1 expression vector (GenScript Biotech, Netherlands) producing a recombinant protein which contains an N-terminal hexahistidine and Kanamycin resistance. All the recombinant proteins were transformed and expressed in *E*.*coli* BL21 (DE3) (Invitrogen, USA).

Growth of Keio *ΔproB* knockouts complemented with glutamine synthetase variants Complementation assays were performed to ensure that the glutamine synthetase variants actually rescued the Keio ΔproB strain. For this purpose, the knockout strain was transformed by electroporation with each of the Kpn1/Xho1-digested pET-45b(+) expression vectors (GenScript Biotech, Netherlands), containing the single glutamine synthetase variants and selected by addition of 50 μg/mL kanamycin and 100 μg/mL ampicillin to the minimal medium. All cultures were grown in minimal medium until saturation (about 36h at 37°C and shaking). They were then diluted to OD_600_= 0.1 and the optical density (OD_600_) of all cultures was measured after 14h (Extended Data Fig. 3).

### Protein purification

Cells harboring the plasmids containing γ-glutamyl phosphate reductase gene or glutamine synthetase genes were grown at 37°C to an OD_600_ of ∼0.6 in Luria-Bertani medium supplemented with 1μg ·mL ^-1^ kanamycin or 100μg ·ml^-1^ ampicillin, respectively. Overexpression of all the recombinant proteins were induced with 0.4 μM isopropyl β-D-thiogalactopyranoside (IPTG) and cell growth was continued at 25°C overnight. The cells were harvested by centrifugation at 6198 g for 15 minutes at 4°C and the pellets were immediately stored at −80°C after their resuspension in cold lysis buffer [20 mM NaPO_4_ pH 7.4 / 0.5 M NaCl / 20mM Imidazole] supplemented with protease inhibitors. Cells were thawed and disrupted by sonication on ice. The insoluble fractions were removed by centrifugation (27,000xg, 30 min at 4°C). Hexahistidine-tagged proteins in the soluble fraction were affinity purified using a nickel-charged His-trap immobilized metal affinity chromatography (IMAC) column (GE Healthcare, UK). Washing was carried out with lysis buffer and recombinant proteins were eluted with the same buffer containing 500 mM imidazole. The eluted glutamine synthetase proteins (wild type or single-mutant variants) were then dialyzed against buffer 50 mM Tris–HCl, pH 7.0 / 2mM MgCl_2_, while γ-glutamyl phosphate reductase was dialyzed against buffer 50 mM Tris–HCl, pH 7.0 / 1mM DTT. Purity was confirmed by SDS-PAGE and concentrations were determined spectrophotometrically using molar extinction coefficients at 280 nm of 36580 M^−1^·cm^−1^ for glutamine synthetase variants and 17,795 M^−1^·cm^−1^ for γ-glutamyl phosphate reductase. All proteins were stored at -80°C and at concentrations above 100 μM in the dialysis buffer. No deterioration of the activities was observed upon storing the enzymes at -80 °C, at least over a period of few months.

### Experimental determination of leakage of γ-glutamyl phosphate from glutamine synthetase variants

Since γ-glutamyl phosphate is a highly labile species, leakage was determined through the detection of the cyclization product, pyroglutamic acid. Samples were prepared in 50 mM Tris–HCl, 15 mM ATP, 4 or 3 mM L-glutamate, 25 mM NH_4_Cl, 50 mM MgCl_2_ at pH 7.0. The reaction of glutamine synthetase was started with the addition of 300μg of the enzyme (either wild type of single-mutant variant) to a final volume of 1mL. The solutions were kept at 25°C for 12 hours and their chemical composition was assessed by Mass Spectrometry or Nuclear Magnetic Resonance, as described below.

### Mass Spectrometry

Samples for mass spectrometry were prepared by dilution 1:2000 in water/acetonitrile 40/60 (v/v) and 5.0μL aliquots were injected in an ultraperformance liquid chromatography (UPLC) system consisting of a Quaternary Solvent Manager Acquity (Waters Corporation) equipped with a column Cortecs UPLC Hillic (1.6 μM, 2.1 × 50 mm) and coupled with a triple quadrupole mass spectrometer XEVO-TQS (Waters Corporation). The mobile phase of the UPLC system consisted of H_2_O + 0.1% formic acid v/v (Solvent A) and acetonitrile + 0.1% formic acid v/v (Solvent B). Linear gradient elution (flow rate 0.2mL/min) was programmed as follows: 0-4 min (20% A-80%B), 4-5 min (40% A-60%B), 5-5.1 min (20% A-80%B). Electrospray ionization mass spectra (ESI-MS) were acquired in the positive (ESI +). LC-MS/MS acquisition parameters for the target molecules were as follows: L-Glutamic acid precursor ion 148.0, product ions 84.0 and 101.9; L-Glutamine precursor ion 147.0, product ions 84.0 and 130.0; L-Pyroglutamic acid precursor ion 130.0, product ions 56.0 and 84.0. For each glutamine synthetase variant, three independent determinations were performed.

### Mass spectrometry determination of pyroglutamic acid *in vivo*

Original rescues were grown at 37°C in kanamycin-supplemented minimal medium until they reached an optical density of 0.7. As a control, the knockout strain Keio *ΔproB* was grown in minimal medium supplemented with kanamycin and 50 μM proline. ^13^C- and ^15^N-labelled glutamic acid (CAS number: 202468-31-3) from Cortecnet was added to all samples to a final concentration of 1mM. Afterwards, all samples were kept at 25°C in shaking for 10 min. Cells were collected by centrifugation at 4,000 rpm at 4 °C for 15 min and the supernatant was discarded. Cells were completely resuspended in 1 mL of water/acetonitrile 40:60 (v/v) and centrifuged at 14,000rpm for 15 min. 5μL of the supernatant of each sample was injected in an ultraperformance liquid chromatography (UPLC) system consisting of a Quaternary Solvent Manager Acquity (Waters Corporation) equipped with a column Cortecs UPLC Hillic (1.6 μM, 2.1 × 50 mm) and coupled with a triple quadrupole mass spectrometer XEVO-TQS (Waters Corporation). The mobile phase of the UPLC system consisted of H_2_O + 0.1% formic acid v/v (Solvent A) and acetonitrile + 0.1% formic acid v/v (Solvent B). Gradient elution (flow rate 0.2mL/min) was programmed as follows: 0-4 min (20% A-80%B), 4-5 min (40% A-60%B), 5-5.1 min (20% A-80%B). Electrospray ionization mass spectra (ESI-MS) were acquired in the positive (ESI +). LC-MS/MS acquisition parameters for the target molecules were as follows: L-Pyroglutamic acid precursor ion 130.0, product ion 84.0, [^13^C_5_ ^15^N] L-Pyroglutamic acid precursor ion 136.0, product ion 89.0. In order to quantify the concentration of each of the target molecules, a standard curve was prepared with the pure compounds without isotopic labelling (Zhang P, 2019). For each original rescue and control experiment, three biological replicates were performed.

### Nuclear Magnetic Resonance Spectroscopy

^1^H NMR spectra were acquired on a Bruker Avance III 600 Hz spectrometer at a proton frequency of 600MHz equipped with a TCIP cryoprobe at 298 K. Samples were supplemented with 5% D_2_O and 0.01 mM DSS (4,4-dimethyl-4-silapentane-1-sulfonic acid). The standard zgesgp pulse sequence from the Bruker library was used with excitation sculpting to allow suppression of the water (Huang & Shaka, 1995). Data was collected with 16,384 points and a spectral width of 16.0242 ppm with 256 scans, 16 dummy scans and an acquisition time of 1.7 s. Data was processed using MestreNova software. All spectra were automatically phased, baseline corrected using a polynomial function and calibrated to the center of the DSS peak. The concentrations of glutamate, glutamine and pyroglutamate were measured by integrating the ^1^H peaks corresponding to gamma protons at ^1^H frequencies of 2.33, 2.43 and 2.41 ppm respectively. For each glutamine synthetase variant, three independent determinations were performed.

### Assay of γ-glutamyl phosphate reductase activity coupled with glutamine synthetase

The coupled activity assay we used to assess the diffusion of γ-glutamyl phosphate from glutamine synthetase to γ-glutamyl phosphate reductase is identical to that described by Smith et al. (1984), except for the source of γ-glutamyl phosphate. Specifically, we used variants of glutamine synthetase as source of γ-glutamyl phosphate, instead of γ-glutamyl kinase. The measurement of the reductase activity was based on the spectrophotometric determination of the oxidation of NADPH. The reaction solution was 50 mM Tris–HCl pH 7.0, 15 mM ATP, 8 mM L-glutamate, 25 mM NH_4_Cl, 50 mM MgCl_2_, 0.15mM NADPH and 7μM γ-glutamyl phosphate reductase. The solution was equilibrated at 25 °C for 5 min, the reaction was initiated by addition of a glutamine synthetase variant to a concentration of 7μM. The enzyme activity was determined using the initial reaction rates determined from changes in A_340_ nm and the molar extinction coefficient of NADPH at that wavelength (6221 M^−1^ ·cm^−1^). Three independent coupled activity assays of γ-glutamyl phosphate reductase were performed with each glutamine synthetase variant as source of γ-glutamyl phosphate.

Computational calculation of the barrier for γ-glutamyl phosphate cyclization Calculations were performed by means of density functional theory (DFT) using the GAUSSIAN09 suite (Frisch et al., 2010). Structures were optimized using the widely used B3LYP functional (Becke et al., 1988; Lee et al., 1988; Becke, 1993) with the Pople’s-based 6–31G* basis set (Fancl et al., 1982). All minima were fully optimized by the gradient technique and assessed by frequency calculations. Enthalpy and Gibbs free energy values were obtained by taking into account zero-point energies, thermal motion, and entropy contribution at standard conditions (temperature of 298.15 K, pressure of 1 atm). A Polarizable Continuum Model (PCM) (Miertus et al., 1981; Tomasi & Persico, 1994) has been taken into account to reproduce the solvent in which reactions and/or physical measurements have been carried out (water).

### Data availability

Raw sequencing data are available in the Sequence Read Archive (SRA) under the PRJNA839218 BioProject accession number.

## Acknowledgements

This work was supported by Human Frontier Science Program Grant RGP0041/2017 (J.M.S.-R. and B.S.), Spanish Ministry of Economy and Competitiveness/FEDER Funds Grant RTI2018-097142-B-100 (J.M.S.-R.), National Aeronautics and Space Administration (NASA) Grant 80NSSC18K1277 (B.S.) and Grant E-BIO-464-UGR-20 (E.M.-C.) from FEDER Funds and Consejería de Economía, Conocimiento, Empresas y Universidad de la Junta de Andalucia. E.A.-L. was a recipient of a postdoctoral fellowship from the regional Andalusian Government (2020_DOC_00541). We thank the “Centro de Supercomputacion” (ALHAMBRA-CSIRC) of the University of Granada for providing computational resources and Dr. Valeria A. Risso for comments on the manuscript.

## Author contributions

E.M.-C. carried out the selection and identification of proline auxotropy rescues, the protein mutagenesis and preparation of protein variants, and the coupled activity assays, under the supervision of B.I.-M., who also provided essential input regarding the experimental design of the study; F.M.-T. and M.S.N. set up the experimental protocol for the determination of proline prototrophy restauration under the supervision of B.S.; L.I.G.-R. carried out preliminary experiments to explore and eventually rule out alternative explanations for proline prototrophy restauration; E.M.-C. and G.G.-A. carried out the NMR experiments under the supervision of J.L.O.-R.; E.M.-C. and M.O.-M. carried out the mass spectrometry experiments; E.A.-L. performed the bioinformatics analyses for genomic DNA sequencing; A.J.M. and P.R. performed the computational calculations on the cyclization barrier under the supervision of J.M.C.; J.M.S.-R. conceived and designed the research, and wrote the first draft of the manuscript; all authors discussed the manuscript, suggested modifications and improvements, and contributed to the final version.

## Competing interests

The authors declare no competing interests

**Correspondence and request for materials:** Jose M. Sanchez-Ruiz

**Extended Data Fig. 1.**
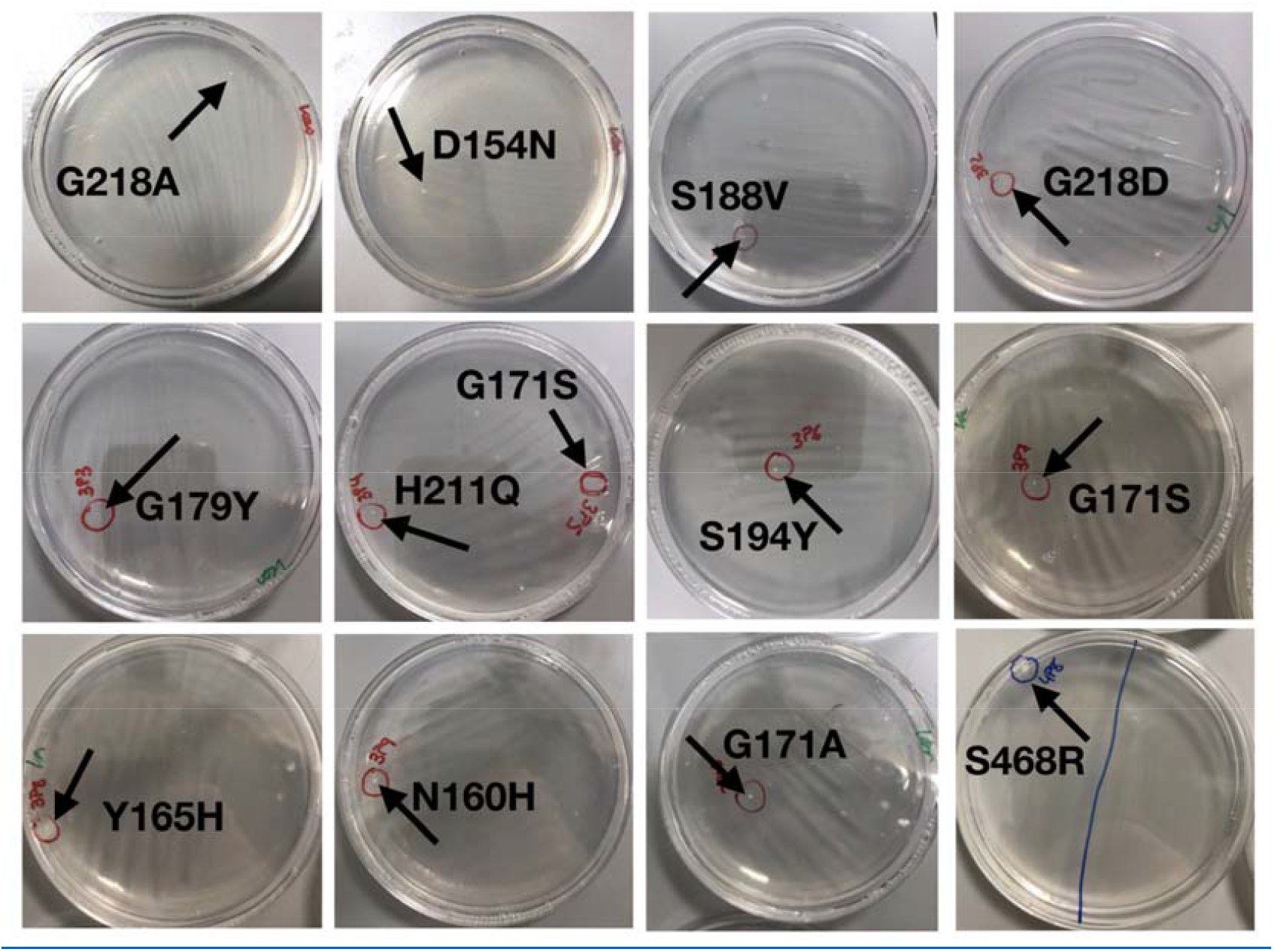
Colonies found upon plating the *ΔproB* knockout strain onto minimal medium. Several hundred plates were examined for four days. We show 12 representative examples of rescues linked to the 12 mutations in the gene of glutamine synthetase extensively analyzed in this work.

**Extended Data Fig. 2.**
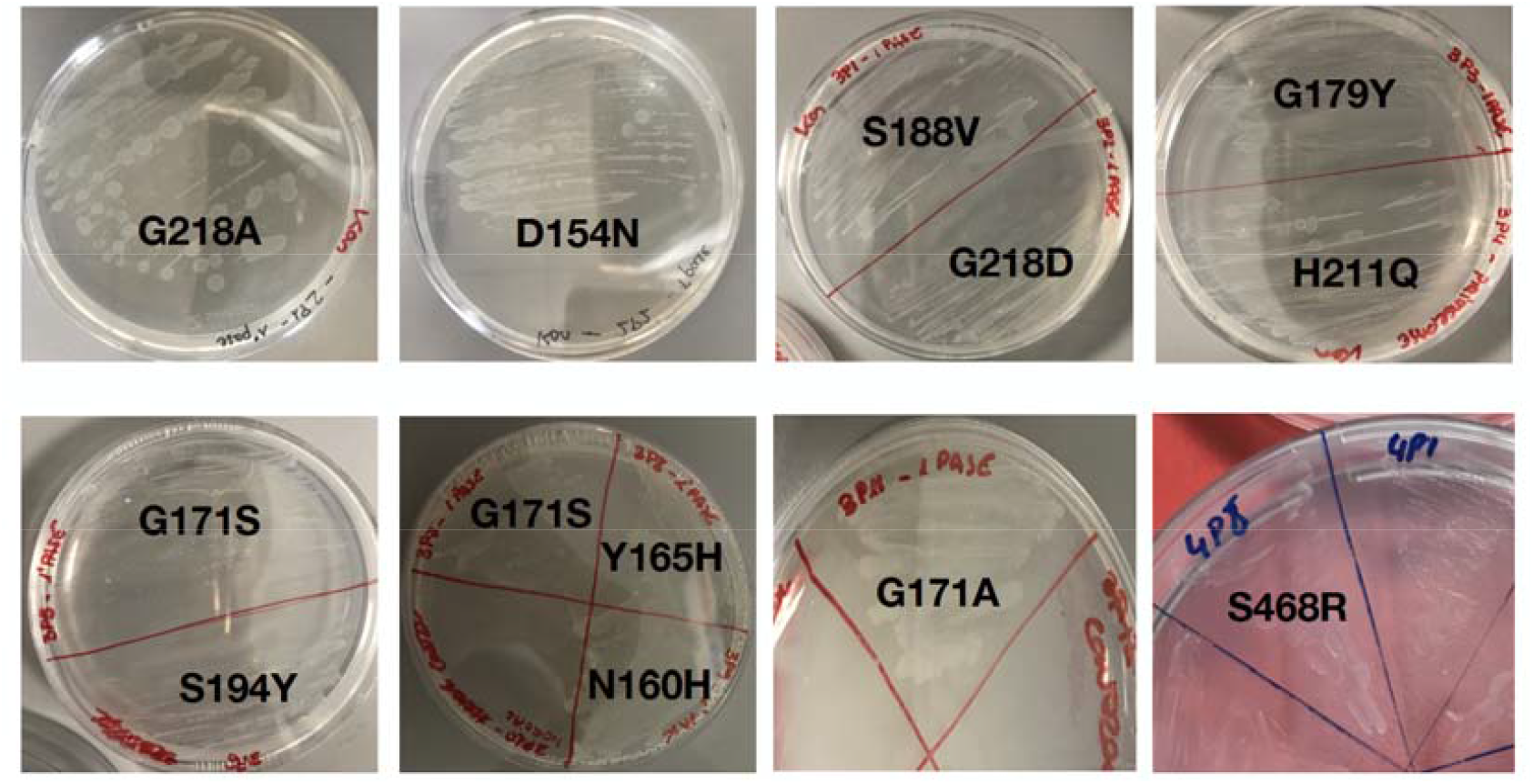
Re-plating onto minimal medium of original rescues. For illustration, only re-plating of representative examples of Extended Data Fig. 1 are shown here, but we found many new colonies upon re-plating in all cases.

**Extended Data Fig. 3.**
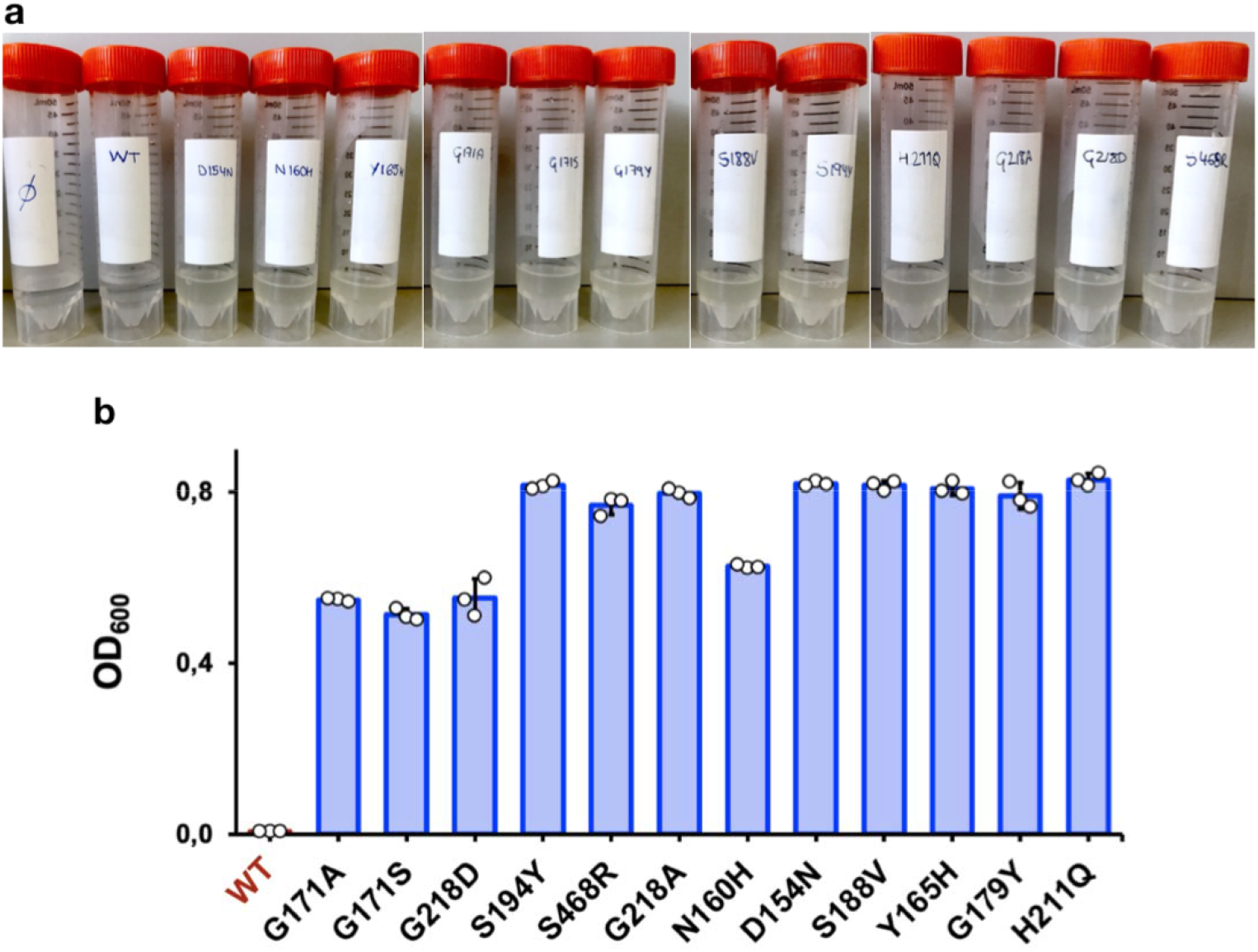
Growth of *ΔproB* knockout cells upon complementation with a plasmid including variants of glutamine synthetase. **(a)** Minimal media containing *ΔproB* knockout cells complemented with plasmid expressing the mutated variants of glutamine synthetase, as well as the wild type enzyme as control. The pictures were taken after allowing the cells to grow for 14 hours at 37 ºC. Experiments were performed in triplicate (only one experiment for each variant is shown here as illustration). Turbidity (revealing cell growth) was observed in all cases with the cells complemented with the single-mutant variants of glutamine synthetase, while the samples containing cells complemented with the wild type glutamine synthetase remained clear. The sample labeled “*ϕ*” is a control in which no cells are present. **(b)** Optical density at 600 nm for the experiments described in a after allowing the cells to grow for 14 hours at 37 °C (see Methods for details). The plot shows average values and standard deviation from the three independent determinations performed for each glutamine synthetase variant. The original values for the three biological replicates are also shown with open circles.

**Extended Data Fig. 4.**
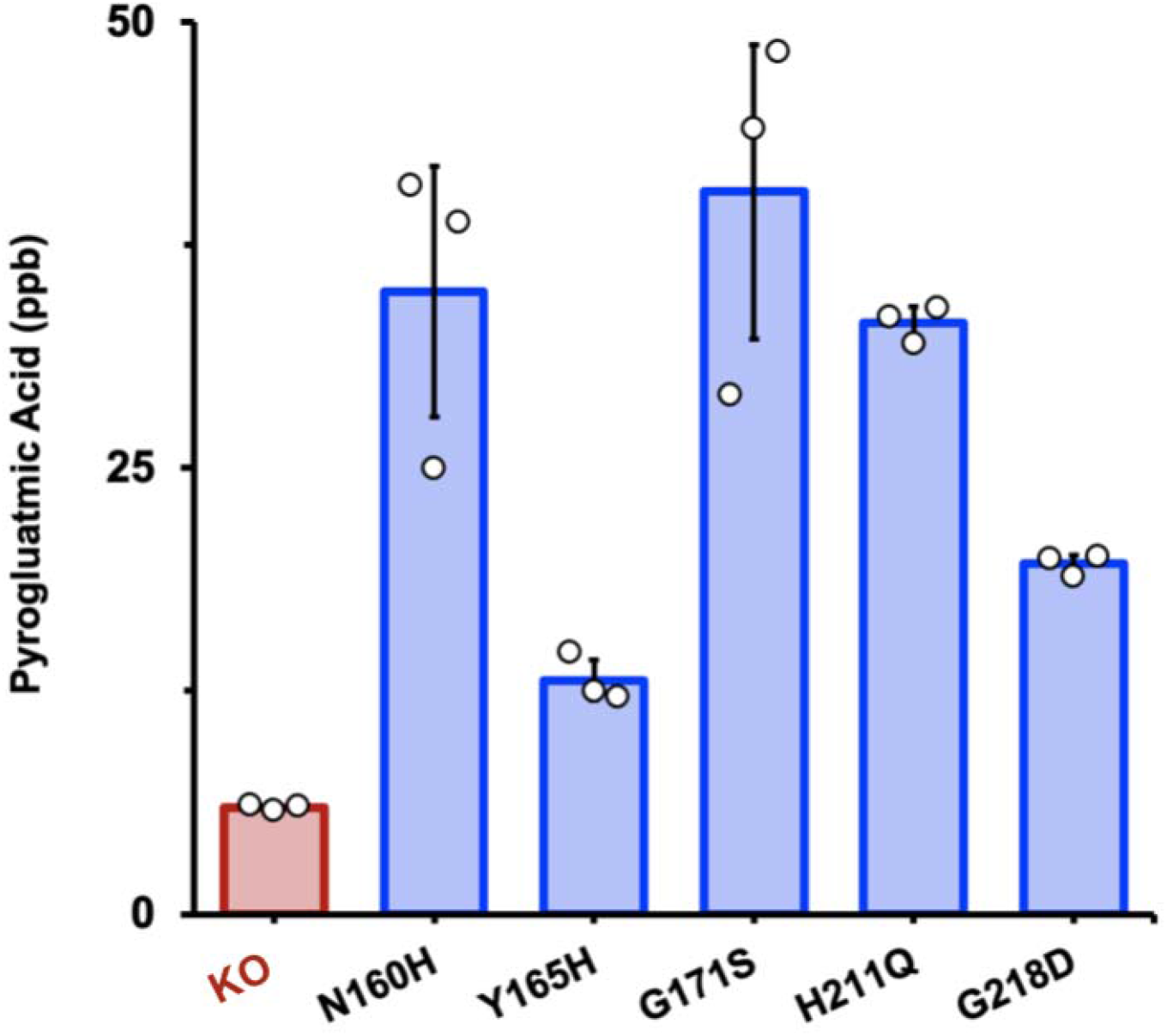
Mass spectrometry determination of pyroglutamic acid *in vivo*. Original rescues of proline auxotrophy were grown in minimal medium, supplemented with isotopically-labeled glutamate and labeled pyroglutamate was determined by mass spectrometry (blue bars). KO refers to a control experiment with the *ΔproB* knockout cells using the same protocol, except that the culture medium was supplemented with proline to enable growth. The values shown are the average of three biological replicates. The original replicate values are shown with open circles and the error bars are standard deviations. The concentrations of pyroglutamic acid shown correspond to the solution obtained after cells separated by centrifugation were completely resuspended in 1 mL of water/acetonitrile 40:60 (v/v) (see Methods for details).

**Extended Data Fig. 5.**
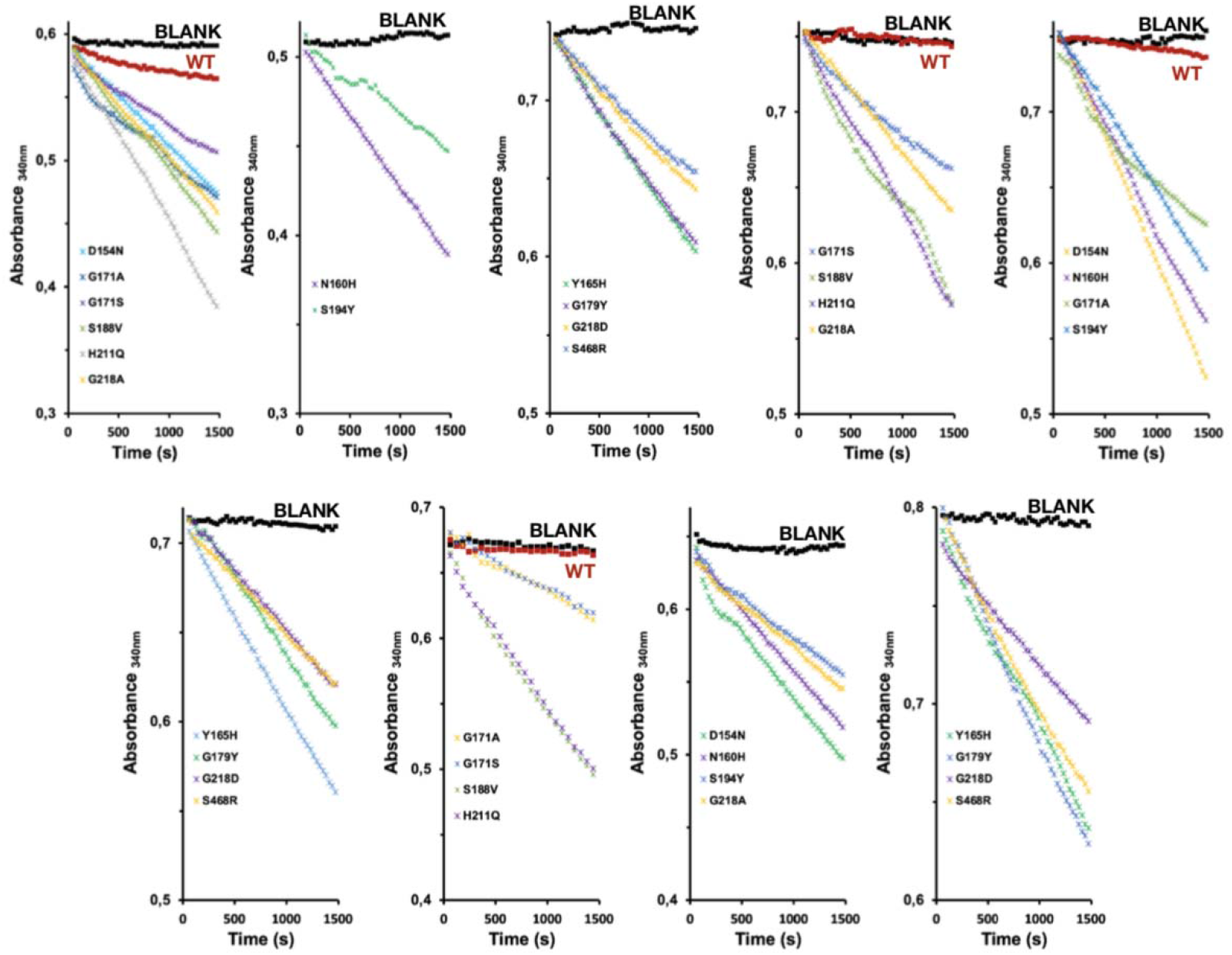
Assay of γ-glutamyl phosphate reductase activity coupled with glutamine synthetase variants. These experiments provided the raw data for the activity values provided in Fig. 4 of the main text. Nine experiments were performed using an UV-VIS spectrophotometer with a multi-cell attachment. In all cases, a control sample (labeled BLANK) in which no glutamine synthetase is present was included. Profiles of absorbance at 340 nm versus time were determined three times with each glutamine synthetase and four times with the wild type protein (profiles labeled WT in red).

**Extended Data Fig. 6.**
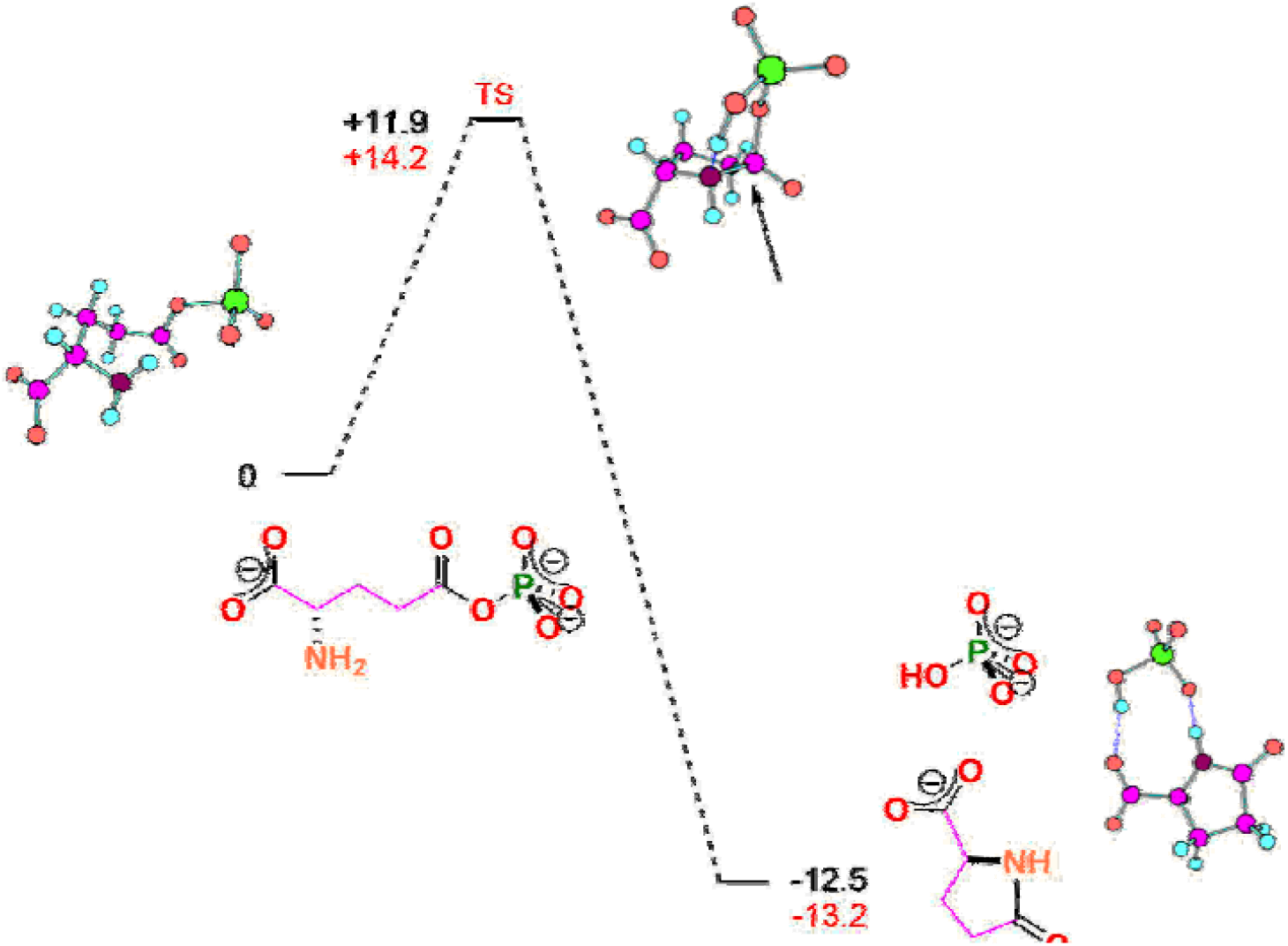
Energy (figures in black) and free energy (figures in red) profiles for the cyclization of γ-glutamyl phosphate with two negative charges at the phosphate group. The profiles have been calculated using density functional theory as implemented in the GAUSSIAN09 suite (see Methods for details). The calculated structures of the reactants, products and the transition state are shown with the following color code for the atoms: carbon (pink), oxygen (red), hydrogen (blue), phosphorus (green) and nitrogen (brown). The reaction involves the nucleophilic intramolecular attack of the amine nitrogen on the carbonyl carbon. The value calculated for activation free energy, 14.2 kcal/mol, leads to a rate constant of 2.4·10^2^ s^-1^ upon application of the Eyring equation from absolute reaction rate theory. However, this value would describe the rate of the reaction when the amine group is fully deprotonated, while, at neutral pH, most of the amine group will be in the protonated form, which is not competent for the reaction. Assuming that the pK value of amine group in γ-glutamyl phosphate is similar to that in glutamic acid (9.7), the fraction of deprotonated amine at pH 7 is estimated to be about 2·10^−3^. Correction for this fraction leads to an estimated rate constant of about 0.5 s^-1^ and a time scale for the reaction of a couple of seconds, which is expected to be orders of magnitude above the time required for a small molecule to diffuse across an *E. coli* cell. For instance, the time for diffusion of a typical protein across an *E. coli* cell has been estimated to be about 10 ms (Philips et al., 2009) and diffusion of a small molecule is expected to be faster. Certainly, the correction for protonation of the amine we have performed (*i*.*e*., multiplying by the fraction of deprotonated amine), implicitly assumes that cyclization is rate determining and that protonation-deprotonation of the amine is fast and remains at equilibrium. It could be conceivable that amine deprotonation could become rate-determining at neutral, but as with any change in rate determining step, this would only decrease the overall rate of the reaction thus increasing the gap between the predicted time scale for cyclization and the time associated to diffusion across an *E. coli* cell.

**Extended Data Fig. 7.**
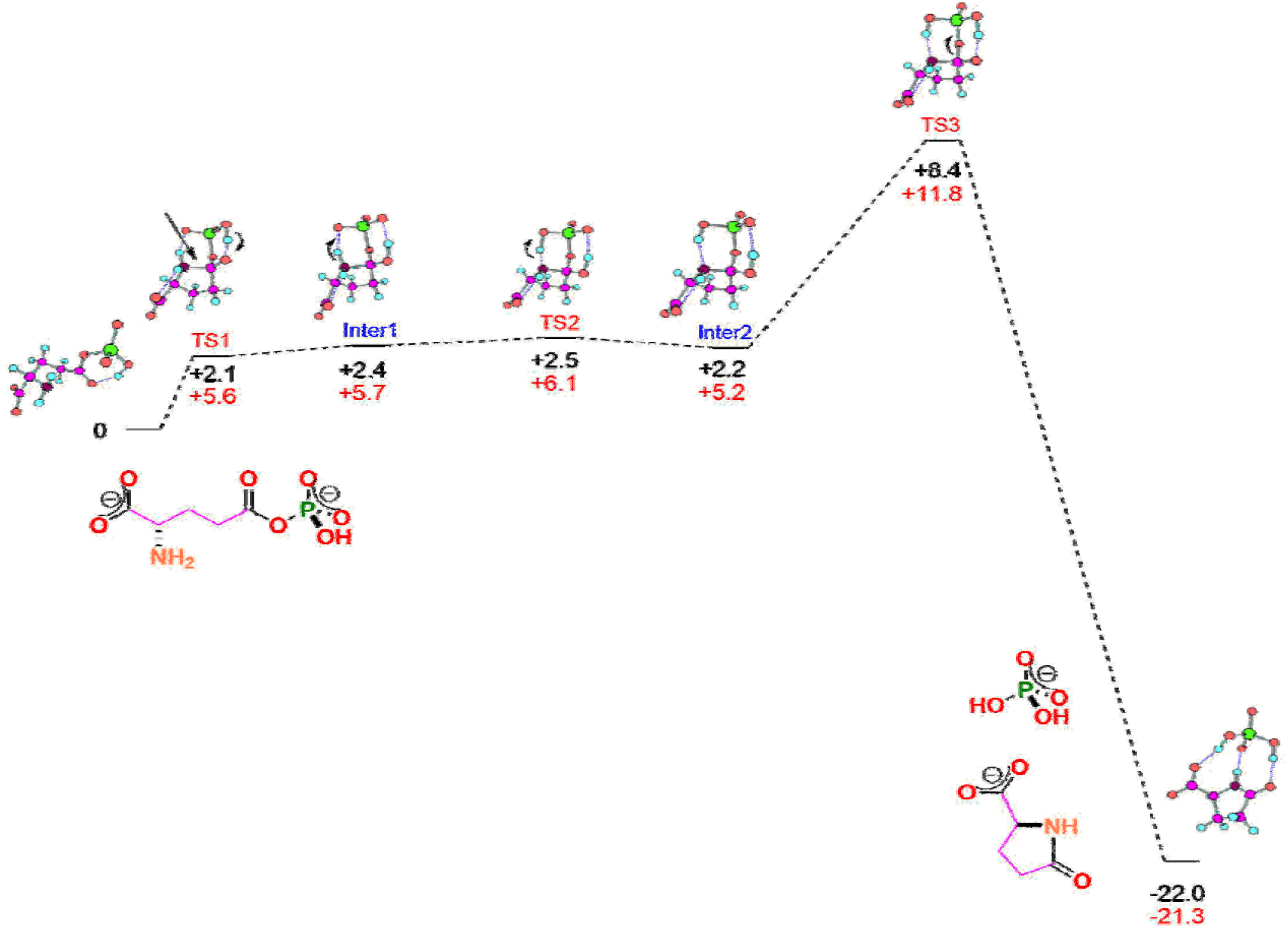
Energy (figures in black) and free energy (figures in red) profiles for the cyclization of γ-glutamyl phosphate with one negative charge at the phosphate group. The profiles have been calculated using density functional theory as implemented in the GAUSSIAN09 suite (see Methods for details). The calculated structures of the reactants, products and the transition state are shown with the following color code for the atoms: carbon (pink), oxygen (red), hydrogen (blue), phosphorus (green) and nitrogen (brown). The reaction involves the nucleophilic intramolecular attack of the amine nitrogen on the carbonyl carbon. However, unlike the cyclization of the γ-glutamyl phosphate with two negative charges at the phosphate group (Extended Data Fig. 4), here proton re-arrangement must occur prior to the efficient release of the phosphate group. The value calculated for activation free energy, 11.8 kcal/mol, leads to a rate constant of 1.4·10^4^ s^-1^ upon application of the Eyring equation from absolute reaction rate theory. However, this value would describe the rate of the reaction when the amine group is fully deprotonated, while, at neutral pH, most of the amine group will be in the protonated form, which is not competent for the reaction. Assuming that the pK value of amine group in γ-glutamyl phosphate is similar to that in glutamic acid (9.7), the fraction of deprotonated amine at pH 7 is estimated to be about 2·10^−3^. Correction for this fraction leads to an estimated rate constant of about 28 s^-1^ and a time scale for the reaction of about 36 ms, which is expected to be clearly above the time required for a small molecule to diffuse across an *E. coli* cell. For instance, the time for diffusion of a typical protein across an *E. coli* cell has been estimated to be about 10 ms (Philips et al., 2009) and diffusion of a small molecule is expected to be faster. Certainly, the correction for protonation of the amine we have performed (i.e., multiplying by the fraction of deprotonated amine), implicitly assumes that cyclization is rate determining and that protonation-deprotonation of the amine is fast and remains at equilibrium. It could be conceivable that amine deprotonation could become rate-determining at neutral, but as with any change in rate determining step, this would only decrease the overall rate of the reaction thus increasing the gap between the predicted time scale for cyclization and the time associated to diffusion across an *E. coli* cell.

**Extended Data Fig. 8.**
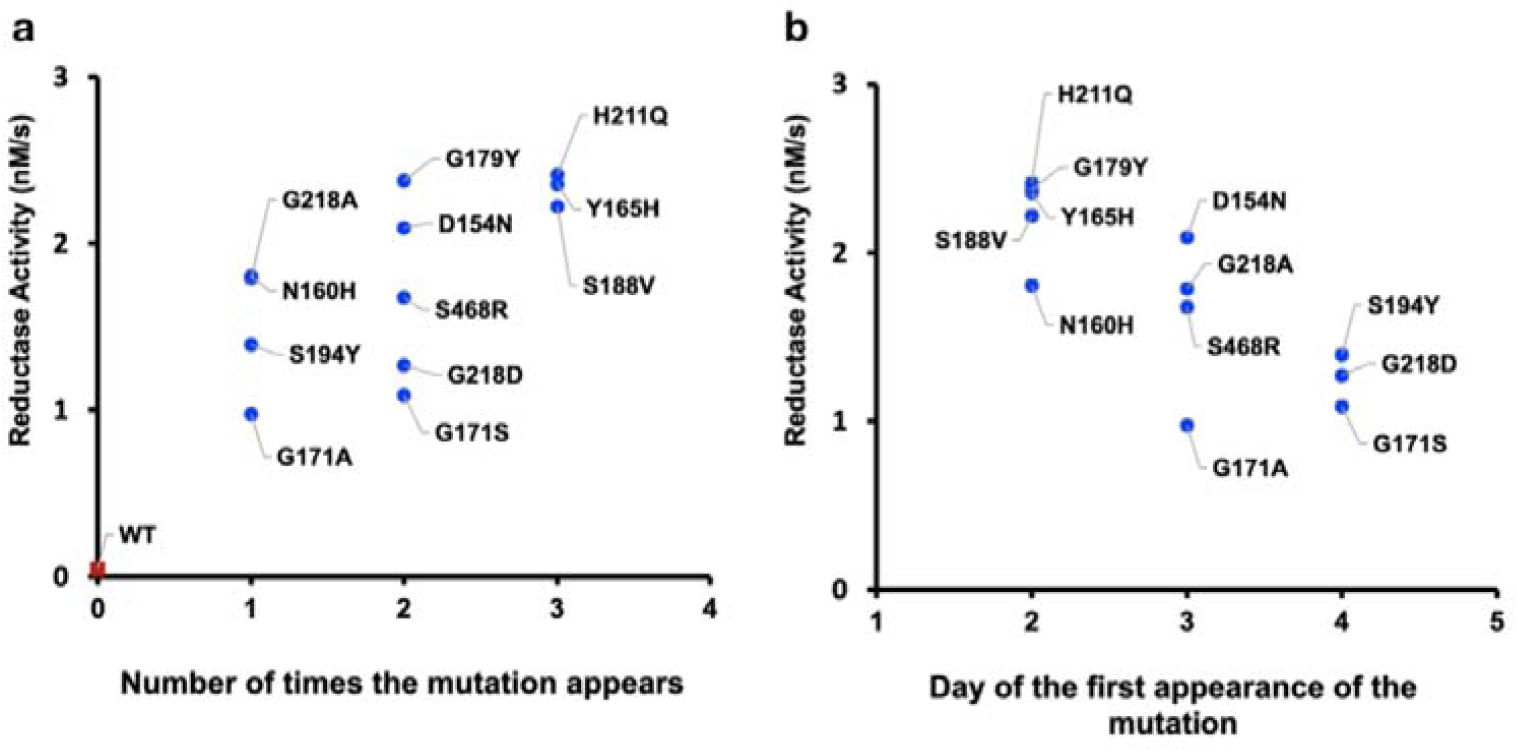
γ-glutamyl phosphate reductase activity in the coupled assay with glutamine synthetase variants (Fig. 4) versus efficiency of restoration of proline prototrophy. Two metrics of restoration efficiency are used, the number of times the mutation appears after plating of the knockout strain onto minimal medium (a) or the day of the first appearance of the mutation (b). The plot of activity versus the number of times the mutation appears (a) has a correlation coefficient of 0.77 and a p value (probability that the correlation arises by chance) of 2·10^−3^. The plot of activity versus the day of the first appearance of the mutation (b) has a correlation coefficient of 0.82 and a p value (probability that the correlation arises by chance) of 10^−2^.

## References

1. Almassy, R.J., Janson, C.A., Hamlin, R., Xuong, N.-H. & Eisenberg, D. Novel subunit-subunit interactions in the structure of glutamine synthetase. Nature 323, 304–309 (1986).

2. Baba, T., Ara, T., Hasegawa, M., Takau, Y., Okumura, Y., Baba, M., Datsenko, K.A, Tomita, M. Wanner, B.L. & Mori, H. Construction of Escherichia coli K-12 in-frame, single-gene knockout mutants: the Keio collection. Mol. Syst. Biol. 2, 2006.0008 (2006).

3. Berg, C.M. & Rossi, J.J. Proline excretion and indirect suppression in Escherichia coli and Salmonella typhimurium. J. Bacteriol. 118, 928–938 (1974).

4. Copley, S.D. Evolution of efficient pathways for degradation of anthropogenic chemicals. Nat. Chem. Biol. 5, 559–566 (2009).

5. Copley, S.D. An evolutionary biochemist’s perspective on promiscuity. Trends Biochem. Sci. 40, 72–78 (2015).

6. Cravens, A., Payne, J. & Smolke, C.D. Synthetic biology strategies for microbial biosynthesis of plant natural products. Nat. Comunn. 10, 2142 (2019).

7. De Crécy-Lagard, V., Haas, D. & Hanson, A.D. Newly-discovered enzymes that function in metabolite damage-control. Curr. Opin. Chem. Biol. 47, 101–108 (2018).

8. Csonka, L.N. & Leisinger, T. Biosynhesis of proline. EcoSalPlus 2013; doi:10.1128/ecosalplus.3.6.1.4

9. Digianantonio, K.M. & Hetch, M.H. A protein constructed de novo enables cell growth by altering gene regulation. Proc. Natl. Acad. Sci. U.S.A. 113, 2400–2405 (2016).

10. Digianantonio, K.M., Korolev, M. & Hetch, M.H. A non-natural protein rescues cells deleted for a key enzyme in central metabolism. ACS Synth. Biol. 6, 694–700 (2017).

11. Eisenberg, D., Gill, H.S., Pfluegl, G.M.U. & Rotstein. S.H. Structure-function relationships of glutamine synthetases. Biochim. Biophys. Acta 1477, 122–145 (2000).

12. Fichman, Y., Gerdes, S.Y., Kovács, H., Szabados, L., Zilberstein, A. & Csonka, L.N. Evolution of proline biosynthesis: enzymology, bioinformatics, genetics, and transcriptional regulation. Biol. Rev. 90, 1065–1099 (2015).

13. Grunwald, P. Biocatalysis. 2nd ed. World Scientific, Singapore (2018).

14. Huang, X.H., Holden, H.M. & Raushel, F.M. Channeling of substrates and intermediates in enzyme-catalyzed reactions. Annu. Rev. Biochem. 70, 149–180 (2001).

15. Khersonsky O. & Tawfik, D.S. Enzyme Promiscuity: a mechanistic and evolutionary perspective. Annu. Rev. Biochem. 79, 471–505 (2010).

16. Kim, J., Kershner, J.P., Novikov, Y., Shoemaker, R.K. & Copley, S.D. The serendipitious pathways in E. coli can bypass a block in pyridoxal-5’-phosphate synthesis. Mol. Syst. Biol. 6, 436 (2010).

17. Kim J., Flood, J.J., Kristofich, M.R., Gidfar, C., Morgenthaler, A.B., Fuhrer, T., Sauer, U., Snyder, D., Cooper, V.S., Ebmeier, C.C., Old, W.M. & Copley, S.D. Hidden resources in the Escherichia coli genome restore PLP synthesis and robust growth after the deletion of the essential gene pdxB. Proc. Natl. Acad. Sci. U.S.A. 116, 24164–24173 (2019).

18. Kumar, A. & Bachhawat, A.K. Pyroglutamic acid: throwing light on a lightly studied metabolite. Curr. Sci. 102, 288–297 (2012).

19. Lee, H., Phuong, N.L. & Na, D. Advancement of metabolic engineering assisted by synthetic biology. Catalysts 8, 619 (2018).

20. Lerma-Ortiz, C., Jeffryes, J.G., Cooper, A.J.L., Niehaus, T.D., Thamm, A.M.K., Frelin, O., Aunins, T., Fiehn, O., de Crécy-Lagard, V., Henry, C.S. & Hanson, A.D. ‘Nothing of chemistry disappears in biology’: the top 30 damage-prone endogenous metabolites. Biochem. Soc. Trans. 44, 961–971 (2016).

21. McLoughlin, S.Y. & Copley, S.D. A compromise required by gene sharing enables survival: implications for evolution of new enzyme activities. Proc. Natl. Acad. Sci. U.S.A. 36, 13497–13502 (2008).

22. Noda-Garcia, L., Liebermeister, W & Tawfik, D.S. Metabolite-enzyme coevolution: from single enzymes to metabolic pathways and networks. Annu. Rev. Biochem. 87, 187–216 (2018).

23. Pál, C., Papp, B. & Lercher, M.J. Adaptive evolution of bacterial metabolic networks by horizontal gene transfer. Nat. Genet. 12, 1372–1375 (2005).

24. Pareek, V., Sha, Z., He, J., Wingreen, N.S. & Benkovic, S.J. Metabolic channeling: predictions, deductions, and evidence. Mol. Cell 16, 3775–3785 (2021).

25. Patrick, W.M., Quandt, E.M., Swartzlander, D.B. & Matsumura, I. Multicopy suppression underpins metabolic evolvability. Mol. Biol. Evol. 24, 2716–2722 (2007).

26. Perachhi, A. The limits of enzyme specificity and the evolution of metabolism. Trends Biochem. Sci. 43, 984–996 (2018).

27. Philips, R., Kondev, J. & Theriot, J. When: stopwatches at many scales. Chapter 3 in “Physical Biology of the Cell”, Garland Science, New York, pp 800 (2009).

28. Schulenburg, C. & Miller, B.G. Enzyme recruitment and its role in metabolic expansion. Biochemistry 53, 836–845 (2014).

29. Srere, P.A. Complexes of sequential metabolic enzymes. Annu, Rev. Biochem. 56, 89–124 (1987).

30. Smith C.J., Deutch, A.H. & Rushlow, K.E. Purification and characteristics of a gamma-glutamyl kinase involved in Escherichia coli proline biosynthesis. J. Bacteriol. 157, 545–551 (1984).

31. Stadtman, E.R. Regulation of glutamine synthetase activity. EcoSal Plus 2013; doi:10.1128/ecosalplus.3.6.1

## Additional references for methods

32. Andrews, S. FastQC: a quality control tool for high throughput sequence data. Avaliable online at: http://www.bioinformatics.babraham.ac.uk/projects/fastqc (2010).

33. Becke, A.D. Density-functional exchange-energy approximation with correct asymptotic behavior. Phys. Rev. A 38, 3098–3100 (1988).

34. Becke, A.D. A new mixing of Hartree–Fock and local density-functional theories. J. Chem. Phys. 98, 5648–5652 (1993).

35. Francl, M.M., Pietro, W.J., Hehre, W.J., Binkley, J.S., DeFrees, D.J., Pople, J.A. & Gordon, M.S. Self-consistent molecular orbital methods. XXIII. A polarization-type basis set for second-row elements. J. Chem. Phys. 77, 3654–3665 (1982).

36. Frisch, M.J., Trucks, G.W., Schlegel, H.B., Scuseria, G.E., Robb, M.A. et al. Gaussian 09, Revision B.01; Gaussian, Inc.: Wallingford, CT (2010).

37. Garrison, E. & Marth, G. Hapoltype-based detection from short-read sequences. arXiv:1207.3907 (2012).

38. Hwang, T.L. & Shaka, A.J. Water suppression that works. Excitation sculpting using arbitrary wave-forms and pulse-field gradient. J. Mag. Res. A 112, 275–279 (1995).

39. Lee, C., Yang, W. & Parr, R.G. Development of the Colle–Salvetti correlation-energy formula into a functional of the electron density. Phys. Rev. B: Condens. Matter Mater. Phys. 37, 785–789 (1988).

40. Li, H. & Durbin, R. Fast and accurate short read alignment with Burrows-Wheeler transform. Bioinformatics 25, 1754–1760 (2009).

41. Li, H., Handsaker, B., Wysoker, A., Fennel, T. Ruan, J., Homer, N., Marth, G., Abecasis, G. & Durbin, R. The sequence alignment/map format and SAMtools. Bioinformatics 25, 2078–2079 (2009).

42. Martin, M. Cutadapt removes adapter sequences from hogh-throughput sequencing reads. EMBnet J. 17.1, 10–12 (2011).

43. Miertuš, S., Scrocco, E. & Tomasi, J. Electrostatic interaction of a solute with a continuum. A direct utilizaion of AB initio molecular potentials for the prevision of solvent effects. J. Chem. Phys. 55, 117–129 (1981).

44. Seemann, T. Snippy-Rapid haploid variant calling and core SNP phylogeny. GitHub. Avaliable at: github.com/tseemann/snyppy/ (2015).

45. Tomasi, J. & Persico, M. Molecular interactions in solution: An overview of methods based on continuous distributions of the solvent. Chem. Rev. 94, 2027– 2094 (1994).

46. Zhang, P., Chan, W., Ang, I.L., Wei, R., Lam, M.M.T., Lei, K.M.K. & Poon, T.C.W. Revisiting Fragmentation Reactions of Protonated α-Amino Acids by High-Resolution Electrospray Ionization Tandem Mass Spectrometry with Collision-Induced Dissociation. Sci. Rep. 23, 6453 (2019).

